# miRSwitch: Detecting microRNA arm shift and switch events

**DOI:** 10.1101/2020.03.25.007351

**Authors:** Fabian Kern, Jeremy Amand, Ilya Senatorov, Alina Isakova, Christina Backes, Eckart Meese, Andreas Keller, Tobias Fehlmann

## Abstract

Arm selection, the preferential expression of a 3′ or 5′ mature microRNA (miRNA), is a highly dynamic and tissue-specific process. Time-dependent expression shifts or switches between the arms are also relevant for human diseases. We present miRSwitch, a web server to facilitate the analysis and interpretation of arm selection events. Our species-independent tool evaluates pre-processed small non-coding RNA sequencing (sncRNA-seq) data, i.e. expression matrices or output files from miRNA quantification tools (miRDeep2, miRMaster, sRNAbench). miRSwitch highlights potential changes in the distribution of mature miRNAs from the same precursor. Group comparisons from one or several user-provided annotations (e.g. disease states) are possible. Results can be dynamically adjusted by choosing from a continuous range of highly specific to very sensitive parameters. Users can compare potential arm shifts in the provided data to a human reference map of pre-computed arm shift frequencies. We created this map from 46 tissues and 30,521 samples. As case studies we present novel arm shift information in a Alzheimer’s disease biomarker data set and from a comparison of tissues in *Homo sapiens* and *Mus musculus*. In summary, miRSwitch offers a broad range of customised arm switch analyses along with comprehensive visualisations, and is freely available at: https://www.ccb.uni-saarland.de/mirswitch/.

## Introduction

The non-coding parts of mammalian genomes play a major role in shaping the gene regulatory landscape [1, 2]. Still, these mechanisms are only understood to a limited extent. Among the many different classes of small or long non-coding RNA elements [3], which modulate the expression of genes on a transcriptional or translational level, microRNAs (miRNAs) seem to play a key role [4, 5]. In the biogenesis of miRNAs, from nascent pri-miRNAs to mature forms, the precursor hairpin molecules are processed by the enzymes DROSHA and DICER [6]. The product of the second cleavage is an approximately 22-nucleotide long RNA duplex structure. Frequently, one arm of the RNA duplex is preferentially accumulated while the other is predominantly degraded [7]. For most hairpins the dominant mature miRNA is assumed to be the functional product. Gene regulation is carried out by the association of the major form with AGO proteins for RNA-induced silencing complex (RISC) formation and successive binding to reverse complementary target sites in mRNAs, mostly within 3’-untranslated regions [8].

High-throughput data from different tissues [9], aging time-points, developmental stages, and physiological conditions [10, 11] indicate, a minor proportion of mature miRNAs from the opposite arm to originate nonetheless. Formerly, these sequences have been denoted as the miR* sequence. Thermodynamic and structural properties have been postulated as drivers for the selection of the dominant arm from the processed duplex [12]. Previous work also demonstrates that arm shifts are specific for tissue types and the distribution of the dominant mature and the miR* sequence can change [13, 14]. For several precursor hairpins, significant quantities of mature miRNAs from both arms are known [15] and both of them are biologically functional. The dominant arm represses translation by means of AGO1 and the miR* sequence by means of AGO2 [16]. Currently, miRNAs are not annotated as miR* and dominant mature form anymore but precisely denoted as the −3p and −5p mature miRNA. Beyond these observations, earlier studies describe the process of selective arm switches in multiple pathological conditions (e.g. [17, 18, 19, 20, 21, 22]). In 2011, Griffith-Jones *et al*. found that arm usage is encoded in the primary miRNA sequence, but outside the mature miRNA duplex, by analysing the miR-100/10 family in different species [16]. The group also provided evidence for functional shifts in insect miRNA evolution [23]. Arm switches have also been correlated to human pathologies such as breast cancer and other severe disorders [18].

Still, a systematic analysis of arm shift or switch events from high-throughput data has not been proposed yet. Here, we introduce miRSwitch, a tool to find differential arm expression from pre-processed high-throughput expression data. In the context of miRSwitch an arm *shift* is a significant enrichment of the 3′ or 5′ arm in one compared to another condition, while still preserving the identity of the dominant form. An arm *switch* denotes a more extreme scenario where in one condition either arm is the dominant and vice versa in the other condition. The supported type of input of miRSwitch ranges from expression matrices over result files from common high-throughput tools such as miRDeep2 [24], miRMaster [25], or sRNAbench [26] to prominent data sources like The Cancer Genome Atlas (TCGA). Users can upload annotation files or manually annotate the samples before running an analysis. To provide even further insights into arm shift events we facilitate a comparison to a background (reference) map of curated arm switch events in *H. sapiens*. We generated this map from 38, 252 human sncRNA-seq data sets comprising 556 reads from 46 different tissues. To demonstrate the functionality of miRSwitch we evaluate an Alzheimer’s disease data set. As second case study, we compare the frequency of arm switches in human and mouse tissues to test the hypothesis whether arm switches might be conserved across species.

## Materials and Methods

### Differential arm expression detection

In a first step, miRNA-precursor pairs of the uploaded expression files are converted to their miRBase v22 identifier using the miRBaseConverter R package [27] (v1.10.0). Then, the precursors are filtered, such that only precursors that have a miRBase v22 identifier and that have two annotated miRNA arms are kept. Next, we remove precursors with no miRNA that have an expression of at least a user specified minimal number of reads. Then we compute the 5′ −3′ ratio difference for every precursor in every sample. Statistical significance between two levels of one annotation variable is calculated with the Wilcoxon-rank sum and for three or more levels using a Kruskal-Wallis test. In all cases, p-values are corrected for multiple hypothesis testing using the Benjamini-Hochberg correction for controlling the false discovery rate. For annotation variables with two levels we also compute the area under the receiver operating characteristic curve (AUC) as effect size. Given a precursor with a 5′ miRNA expression value of *e*5 and a 3′ miRNA expression value of *e*3, we define *e* = *max*(*e*5, *e*3) and 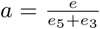 as arm ratio. Also, let *R* be the minimum arm ratio threshold and *X* be the minimum miRNA reads threshold. Precursors are classified as 5′ dominant in one sample, if *a* ≥*R* and *e* ≥*X* and *e*_5_ *> e*_3_, 3′ dominant if *a* ≥*R* and *e* ≥*X* and *e*_5_ *< e*_3_, not dominant if *a < R* and *e* ≥*X*, and not expressed if *e < X*. Arm switch candidates can be queried by defining an additional threshold *S*, requiring 5′ and 3′ dominant precursors in at least *S* samples.

### Web server implementation

We implemented miRSwitch using a dockerized Django Web Framework (v2.2) with a MonetDB database backend (v11.35.19). As job scheduler we used the celery software (v4.3.0). To build a user frontend we used Webpack (v4.41.2) in combination with React JS (v16.12.0), Dev Extreme React Grid (v2.3.2), fornac (v1.1.0), Plotly (v1.51.3), and Highcharts (v7.2.1). The specificity and sensitivity trade-off for potential arm shift / switch candidates can be controlled by adjusting three parameters. These are the minimum arm shift ratio *R*, the minimal number of samples where the threshold *R* needs to be exceeded and the minimal number of reads of one miRNA arm required to compute the 5′ to 3′ ratio.

### Human reference map of differential arm expression

To generate a reference map of human arm shift / switch events we collected 38, 252 human sncRNA-seq samples. The data set was compiled from three different sources, the Sequence Read Archive (SRA) [28] (16, 415), TCGA [29] (10, 999), and samples that were made accessible by anonymous users of miRMaster [25] and who provided consent for aggregated secondary usage (10, 838). Subsequently, we removed duplicated data sets that occurred after pooling the SRA and miRMaster samples. All samples were processed as previously described [30]. Briefly, samples were mapped against the human genome (hg38) and discarded from further analyses in case less than 50% of reads could be aligned with Bowtie (v1.1.2) [31] while allowing no mismatches. We also discarded samples for which either at least 1% of reads mapped to coding regions or fewer than 1 million total reads were detected.

After applying all filtering steps, 30,521 samples remained for consideration. For the samples from SRA, TCGA, and a subset of the miRMaster samples for which annotations were known, the annotation metadata was included in the web server. Other samples for which no tissue annotation was available were labelled as “Unknown”.

### Case studies

To demonstrate the functionality of miRSwitch we performed two case studies; a human liquid biopsy biomarker study and a consideration of arm switches in human compared to mouse tissues. Previously, we obtained sncRNA-seq data from blood of Alzheimer’s disease patients and controls, and validated the data using RT-qPCR [32, 33, 34]. For 70 samples, reads were mapped and miRNA counts for miRBase v21 entries were quantified using miRDeep2. This case study is linked as example data set on the miRSwitch analysis page. As second case study we investigate arm switch events of the same tissues between mouse and human.

## Results

In its essence, miRSwitch allows to search for differential miRNA arm expression between any kind of biological condition. In the following, the basic workflow as well as details and examples on the individual steps are described. The results are concluded with two case studies to demonstrate the features on real-world scenarios where new biological insights can be inferred.

### Workflow of miRSwitch

The workflow of a miRSwitch analysis entails four steps, (1) the input specification; (2) the main analysis functionality; (3) an optional comparisons to the human reference map; and (4) the representation and export of results (Figure 1). Each of the steps is described below in more detail.

**Figure 1:**
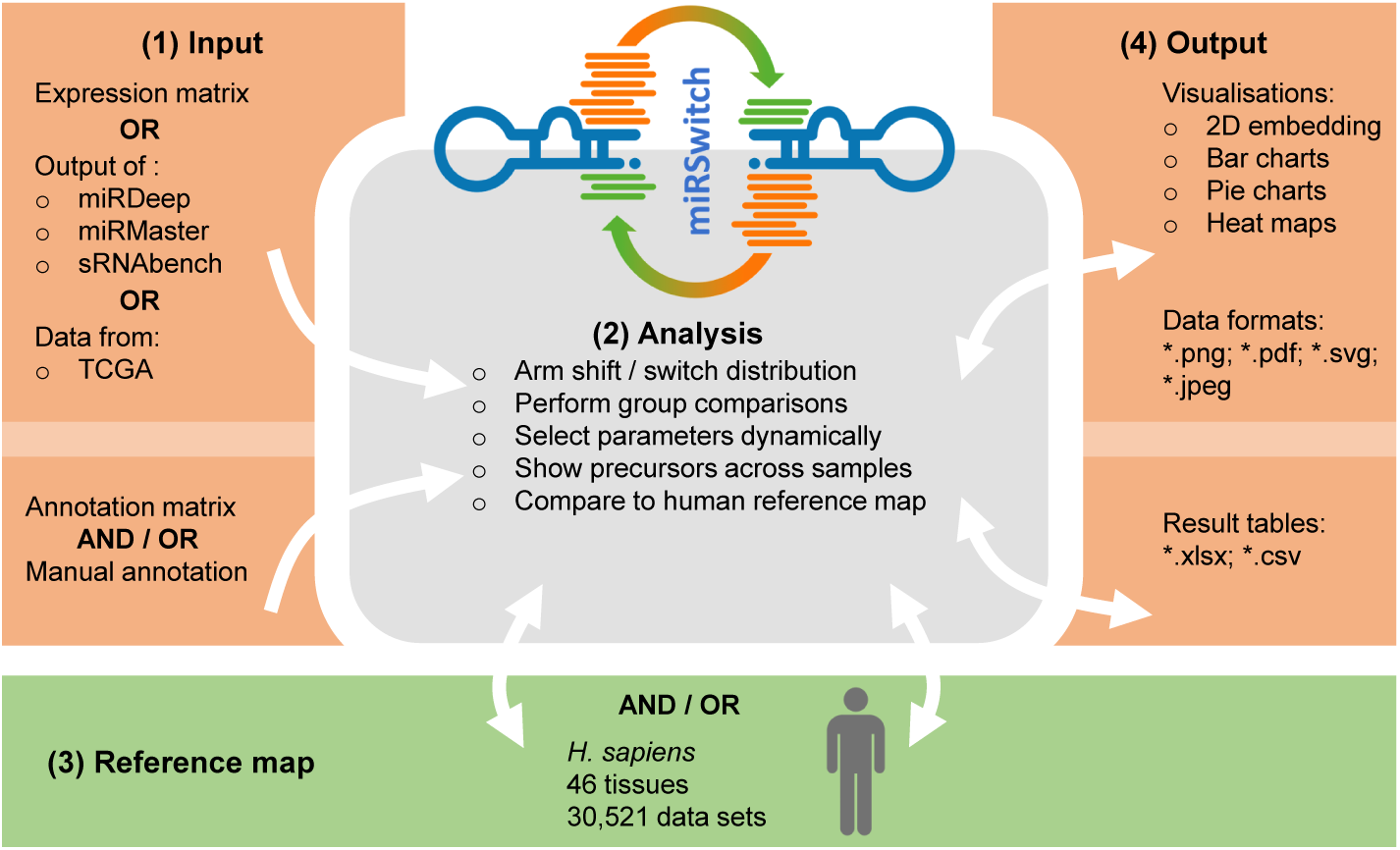
The miRSwitch workflow. miRSwitch consists of four steps (modules), (1) input, (2) miRSwitch analysis core, (3) reference map and (4) output. The arrows denote possible interactions between those. From the input module data are transferred unidirectional to the core module. Between the core and the background map as well as results module bidirectional communication allows to adjust the results dynamically if parameters to define arm switches are changed by the user. All obtained results are easily exportable in common data table formats and plot graphics.

### Step 1: Input specification

As the first step, miRSwitch provides a very flexible user interface to transfer data to the server. For example, the user provides miRNA expression data and an annotation matrix file (.csv, .tsv). The expression data can be uploaded either as a expression matrix file or the plain output from common miRNA discovery and quantification tools applied up-stream. The supported pre-processing tools include miRDeep2, miRMaster, and sRNAbench. Unmodified data files from TCGA can be uploaded as well. Next, the interface automatically extracts metadata from the uploaded file(s) to be displayed in an interactive sample table. Finally, the user can check and modify any derived annotations. In any case, annotation files are optional and annotation variables can be entered manually instead. Most importantly, the first column is expected to match the sample IDs exactly.

### Step 2: Arm expression analysis

From an expression matrix, 5′ and 3′ ratios are computed for each miRNA precursor with two mature forms mapped. Also, the main parameters defining the strength and frequency of arm shift / switch events can be modified. These include the threshold ratio *R* of the 5′ or 3′ arm, the number of reads that have to match to at least one corresponding mature form for each observation and the number of samples that show a respective ratio. Interactive charts and a table of precursors and their classification with respect to the arm dominance are shown on the general analysis page.

Given sample annotations, miRSwitch also performs group comparisons. The user can access the respective results from the “Annotation” tab (for convenience reasons the tab is always named according to the information provided by the user) on the main results page. This tab presents significance values and the area under the receiver operator characteristics curve (AUC) for the group comparisons, as well as a UMAP and PCA embedding of the samples, highlighted according to their group. miRSwitch then computes graphical representations and spreadsheets for all results (see module 4). Since the parameter choice is overall crucial and different researchers may want to use their own specific or sensitive parameters to determine whether a miRNA is able to perform an arm switch, we enable real-time parameter selection and filtering of the respective miRNAs in the web interface.

### Step 3: Reference map

To provide further insights into arm shifts in *H. sapiens* we integrated a reference map of arm shift events in different solid tissues and bio fluids. We collected 38, 252 sncRNA data sets from three different sources (the sequence read archive [28], the cancer genome atlas [29] and data collected by our tool miRMaster [25]). These data sets contain a total of 556 billion sequencing reads and can be annotated for 46 different tissue types. After a stringent quality filtering to exclude data with almost no reads mapping to *Homo sapiens* or containing other sequences but not sncRNAs, a total of 30,521 remained.

Three scenarios from the human reference map demonstrate the importance of the main analysis parameters (cf. Materials and Methods & Results, Step 2). First, to obtain a very specific view, we set the arm ratio of 3′ or 5′ to be at least 80%, at least one miRNA needs to express 1, 000 reads and at least 200 experiments have to show a dominant 5′ expression and another 200 a dominant 3′ expression for the mature miRNAs. For this parameter set (80, 1000, 200), 52 precursors that perform an arm switch are identified, with the most variable being hsa-mir-193a, hsa-mir-30e, hsa-let-7d, hsa-mir-144, hsa-mir-361, and hsa-mir-423. If we alter the parameter set to be less specific (70, 500, 100), already 108 precursors with potential arm switches are identified. Finally, we test a very sensitive parameter set (60, 200, 20). Here, miRSwitch reports 256 precursors with potential arm switches.

### Step 4: Results representation and export

The fourth module is the representation of results generated by the analysis step (2) or extracted from the reference map in step (3). Generally, three different types of results pages are available. Two of them for user provided input and one for the background map. Additionally, detailed results for individual precursors can be viewed in both modules. If an own data set is evaluated the first tab covers general aspects. Aggregated information on how many samples and which annotations were processed, how many precursors were expressed, and how many reads per sample were available. Bubble plots represent how many miRNAs per annotation on the two arms were found and the arm distribution across samples is shown as heat map. Following this general information the user can adjust the parameters for defining arm shift events. The following bar graphs and tables are adjusted dynamically. Here, users can filter or sort the precursors. From the table, single precursors can be selected and detailed information for these candidates are provided. These include links to external databases (miRBase [35] and miRCarta [15]), the sequence and structure, a pie chart on the arm distribution and detailed distribution per annotation group. All information can be displayed either as percentages or absolute values in the bar diagrams. Finally, the information in which samples the miRNA precursor was expressed on both arms is available as interactive table. Next, the annotation tab contains more details about the comparisons of annotation variables. First, sample embeddings from Uniform Manifold Approximation and Projection (UMAP) and Principal Components Analysis (PCA) [37] of the 5′−3′ ratio matrix are provided. Here, color and shape of the points represent the annotation levels for the respective samples. Then, P-values for the difference of the 3′ to 5′ ratio are computed and provided as raw- and adjusted significance values. Another output in the interactive result table that allows sorting and filtering is the AUC value. For each miRNA, the 3′ and 5′ distribution per annotation group available as split violin plot.

Finally, the human reference tab contains all data collected in the background map. The representation is similar to the general results obtained after starting a custom arm ratio analysis. This facilitates an easy comparison. For each precursor, the 5′ and 3′ miRNA expression overall and per tissue are shown. The detailed information for each miRNA precursor is then identical to the results provided by the analysis of own data.

Moreover, distribution plots, pie charts and bar charts are computed to get insights into the distribution of the two mature arms of detected precursors. Figure 2 presents a typical example for a 5′ dominant miRNA (mir-142) and a 3′ dominant miRNA (mir-144).

**Figure 2:**
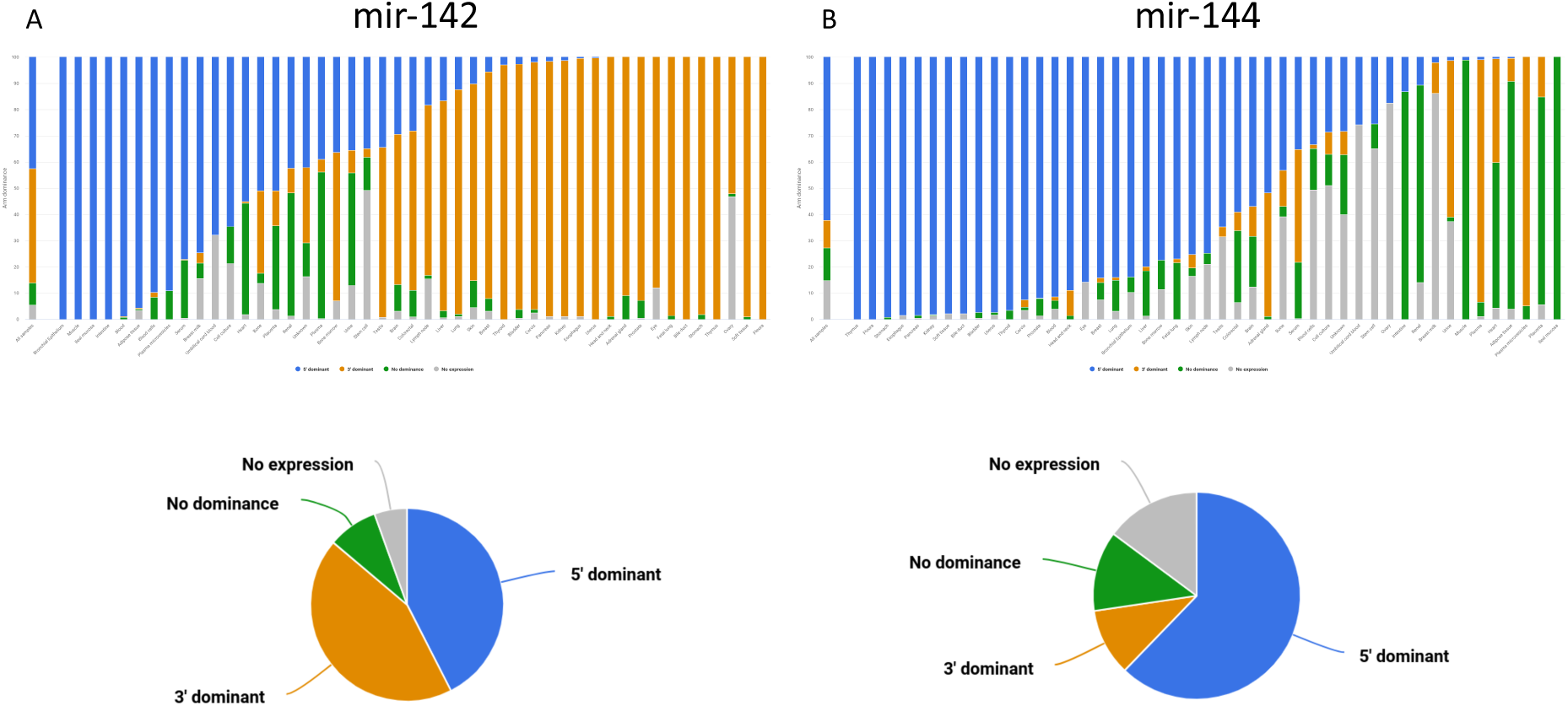
Example of results from the human samples in the background map of arm shift and arm switch events. **(a)** For hsa-mir-142 the distribution of 3′ dominant, 5′ dominant, no dominance, and not expressed are presented for all samples as bar and pie chart. The same classification is also presented for each tissue. For ileal mucosa or blood, the 5′ form is clearly dominant, for thymus or pleura, the 3′ form dominates. **(b)** For hsa-mir-144 the 5′ mature form is clearly more abundant as compared to the 3′ form but in several tissues such as the muscle, no form is dominant.

miRSwitch aims to provide large flexibility with respect to export for down-stream usage. This includes the support of the most relevant image formats covering vector graphics and raster graphics (jpeg, svg, pdf and png). Furthermore detailed result files are also available as spreadsheets. This covers the excel data format as well as the csv plain text file format.

### Case study 1: Alzheimer’s disease

In the first case study we consider previously published sncRNA-seq data from whole-blood samples of Alzheimer’s disease patients and controls. The case study is also provided as example analysis on the miRSwitch homepage. A total of 70 sample files from miRDeep2 are processed in real-time when loading the example and results are available after ∼20 seconds. As first result the web server reports 2, 784 miRNAs and 1, 855 precursors in miRBase v22 of which 748 are expressed with at least one read and have two known mature forms. From the annotation metadata the “Alzheimer” and “Control” labels were identified. Using the default parameter set, our tool reports 10 candidates with arm shift and 3′ dominance as well as 3 candidates with 5′ dominance. Further, the results view indicates that in Alzheimer’s disease fewer 5′ and more 3′ mature miRNAs are expressed as compared to controls (Fig. 3A). The heat map points to one potential outlier sample (SRR837506, data not shown). With the parameter set (80, 5, 3), miRSwitch identifies 7 precursors with 3′ dominant arm (mir-340, mir-199b, mir-29c, mir-6859-1, mir-6859-2, mir-6859-3, and mir-6859-4) and one with a 5′ dominant arm (mir-548h-4). For mir-340, 9% of the samples don’t show an expression, 49% don’t show a dominant arm, 30% are 3′ dominant and 13% are 5′ dominant (Fig. 3B). Dividing this into the two groups of patients and controls we find that except for one sample all samples with 3′ dominant mir-340 come from Alzheimer’s disease patients while control samples are enriched for 5′ dominant mir-340 expression (Fig. 3C). The distribution plot that shows the difference in ratios for controls (left) and patients (right) confirms this on a per sample basis (Fig. 3D). Ultimately, the dodged violin distribution plot demonstrates that in case of mir-340 not only an arm shift but an arm switch between the two groups of samples can be seen (Fig. 3E).

**Figure 3:**
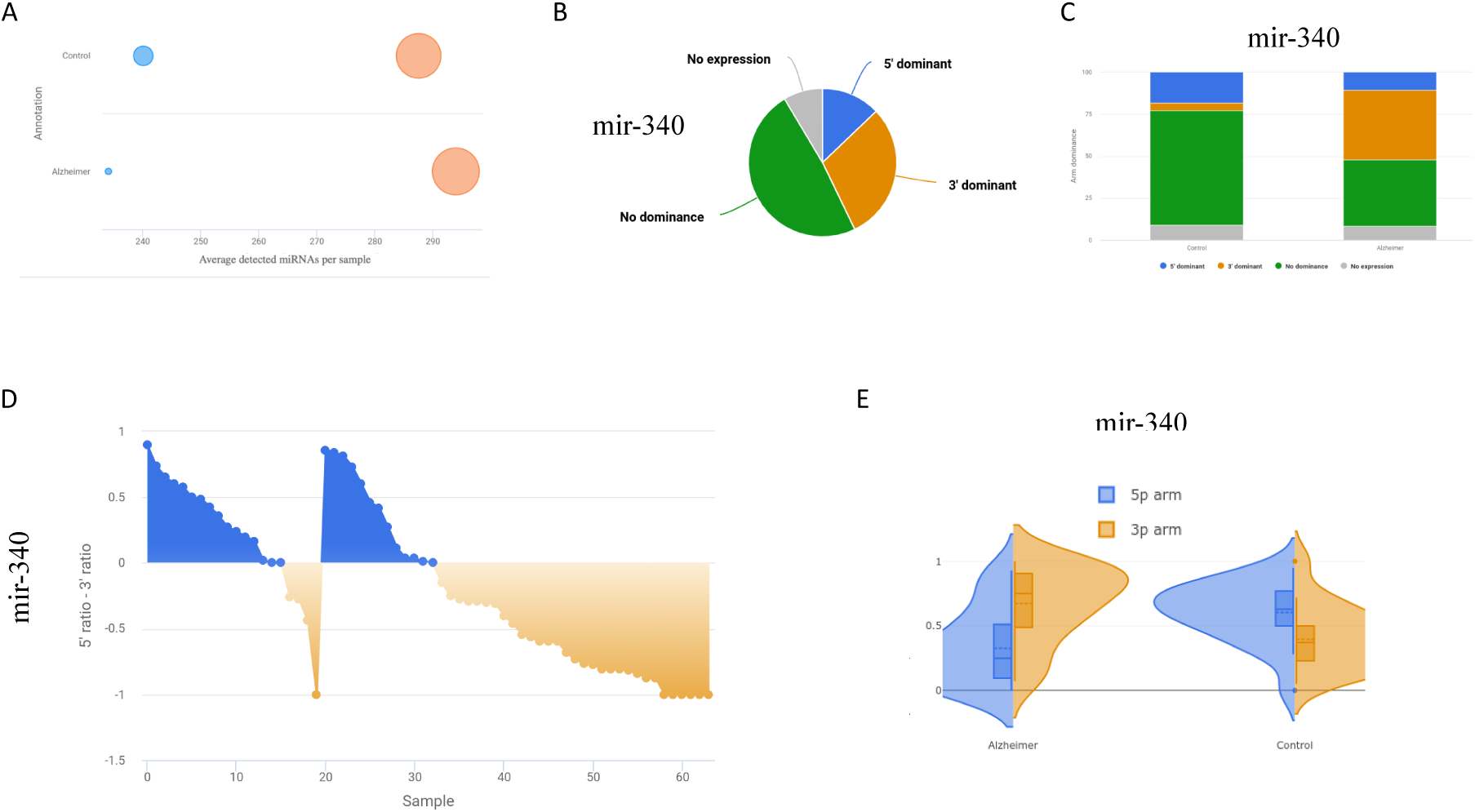
Results of the Alzheimer’s disease case study. **(a)**. Bubble plot showing the number of detected mature miRNAs from both arms for each annotation. **(b)**. Representative pie chart for mir-340 showing in how many samples it is not expressed, shows no arm preference, or is 3′ or 5′ dominant. Plots on the results page are interactive and by hovering over details are displayed. **(c)** Bar chart that splits the information from the pie chart in panel (b) into the annotation levels. **(d)** Distribution chart presenting details on the 5′−3′ difference for the provided groups of samples, here controls (left) and Alzheimer’s disease patients (right). For the latter, a strong enrichment of the 3′ mature form is visible. **(e)** Back-to-back distribution of the arm expression in the provided annotation groups. mir-340 is an arm switch miRNA in Alzheimer’s disease. In the disease it displays higher expression of the 3′ arm while in controls the 5′ arm is more abundant.

### Case study 2: Arm shift events in *M. musculus* and *H. sapiens*

As second case study we analyzed miRNAs from a mouse sncRNA tissue atlas to assess the potential (dis-)similarity of arm shifts in mice and human. To this end, we showcase the value of the human reference map feature to perform the species comparison. We restricted the focus to solid organs included in both data sets.

For example, mir-141 is 3′ dominant in both species, but a higher expression of the 5′ arm was observed for both organisms in testis. Also, mir-26b was largely 5′ dominant in *M. musculus*, only the bone marrow showed expression of both arms. Interestingly, also the human data showed this pattern, although with lower 3′ expression ratios. For mir-106b both organism indicated expression of both arms. In this case, however, a dominant 5′ arm in the human heart was not discovered in mouse samples. mir-337 was mostly 3′ dominant. Brain samples of both organisms however indicate an increased 5′ expression. Although the direct comparison of the mouse tissue data set and the human reference map is biased in its nature, since the latter contains three orders of magnitude more samples from different conditions, we found evidence for many human miRNA arm selection events also in the mice.

## Discussion

Arm shifts and arm switch events have a high impact in many research scenarios, while possible down-stream effects are still underestimated. For example, previous results demonstrate an altered arm distribution between affected and unaffected individuals. Such events have been observed e.g. for breast cancer [18], gastric cancer [22], or prostate cancer [38]. Also the cause of the differences in arm distribution between cell types, developmental stages, and in diseases have been explored only to a limited extent [39, 40]. One likely reason that arm switches have been widely neglected so far, is the missing functionality for arm switch tailored analyses in many standard tools, including our own sncRNA-seq analysis tool miRMaster. The primary goal of this work was to make comprehensive arm switch analyses for any kind of experiment like microarrays, high-throughput sequencing, and RT-qPCR data available to a broader research community. Additionally, well interpretable output is delivered back to the user, both, as interactive graphics and tables. As secondary goal we facilitate the comparison to a reference map containing arm switch events for human. Here, users can check whether their results have already been discovered in any of the previously screened tissues. Remarkably, we consider this resource as a preliminary standardised storage of arm shift events in general, i.e. more human tissues but also other organisms beyond can be added from future experiments.

Proper annotation of miRNAs is still a challenging issue. We used the miRBase V22 [35] annotation. However, not all mature miRNAs might be available in miRBase. In addition, several studies pointed out the erroneous nature of many mature miRNAs in miRBase [15, 41, 42, 30]. Another challenge is the underlying technology. It is well known that expression of miRNAs, similar to other non coding RNAs, mRNAs or proteins, varies depending on the experimental techniques and protocols [15, 41, 42, 43]. Most of the data sets used in the reference map stem from Illumina Sequencing By Synthesis instruments, which may show e.g. a ligation bias [44]. As a consequence, not all potential arm shift events will be discovered due to respective bias. Secondly, the comparison between the user data set and the uploaded data might be compromised and detected differences might occur due to the difference in technologies. To this end, we aim to grow the reference map further, also including other technologies like cPAS Sequencing By Synthesis [43], to promote a less biased comparison and evaluation in the future.

In conclusion, we herein present a very comprehensive web server that facilitates the evaluation of arm shift and arm switch events across different species.

## Data Availability

miRSwitch is freely available at https://www.ccb.uni-saarland.de/mirswitch. No login is required. The data for the case studies are available from the Gene Expression Omnibus (GEO) under accession numbers GSE46579 and GSE119661.

## Supplementary Data

No supplementary data.

## Acknowledgements

We acknowledge the work of the Meese Lab, generating the data set on Alzheimer’s disease.

